# Damage recognition by intestinal stem cells via Draper-Src-Shark-STAT signalling promotes adult *Drosophila* midgut regeneration

**DOI:** 10.1101/2025.03.25.645174

**Authors:** Martina Legido, Rebecca Wafer, Marie Srotyr, Parthive H. Patel

**Affiliations:** School of Cellular and Molecular Medicine, Faculty of Health and Life Sciences, University of Bristol, University Walk, Bristol BS8 1TD UK

## Abstract

Regeneration after injury is crucial for maintaining epithelial structure and function. In order to facilitate this, a regenerative niche forms upon damage that produces cues to induce stem cell proliferation to replace damaged or lost cells. Whereas much is known about this process, less is understood about how stem cells themselves sense damage, translate this into their proliferation and shape the regenerative niche. In the adult *Drosophila* midgut, the non-receptor tyrosine kinase Src42A is required in progenitors to promote ISC proliferation after pathogenic bacterial infection. Although STAT is necessary for Src42A-driven ISC proliferation, the mechanism by which Src42A activates STAT in progenitors remains unclear. Here we show that Draper-Src-Shark signalling acts in ISCs to recognise midgut damage and is required for full STAT activation in ISCs to promote their proliferation. Unlike its role in phagocytes, we find that the engulfment receptor Draper in ISCs does not promote the engulfment of dying midgut epithelial cells but rather is required for ISC proliferation only in the presence of apoptotic enterocytes. Moreover, we uncover that Draper-Src-Shark signalling in progenitors regulates STAT activity non-cell autonomously in the visceral muscle, indicating that progenitors can shape the regenerative niche. As both Src and Stat are activated in the mammalian intestinal epithelium after damage and tumour formation, our work likely extends to mammalian organisms and may aid in developing therapies for tissue regeneration, inflammatory diseases and cancer.

## Introduction

A failure to properly maintain or regenerate an epithelium results in the loss of its integrity, architecture and function, and thus also organismal homeostasis ^1, 2^. In many adult epithelia, stem cells divide to replace damaged or lost tissue cell types ^3, 4^. Stem cells are coaxed to proliferate by cues produced by a regenerative niche that forms shortly after injury ^5–7^. This niche can include other epithelial cells as well as other cell types and tissues associated with the organ. While much is known about how the regenerative niche promotes stem cell proliferation in several adult epithelia, we know less about how stem cells sense damage and how they translate this information to their proliferation. Furthermore, even less is understood about how stem cells modulate their niche or microenvironment after tissue damage to support regeneration.

The mammalian intestinal epithelium is maintained by intestinal stem cells (ISCs) that give rise to differentiated absorptive and secretory epithelial cells ^8^. Upon damage to the mammalian intestine, several cues are received by ISCs that stimulate their proliferation, including Epidermal Growth Factor (EGF), cytokines and Wnts ^5, 6^. These cues are produced by a regenerative niche, which includes resident epithelial cells, immune cells, mesenchyme, vasculature and enteric glial cells ^5, 6^. Similar to the mammalian intestine, the adult *Drosophila* midgut epithelium is maintained by ISCs that produce two main epithelial cell types, absorptive enterocytes (ECs) and secretory enteroendocrine cells (EEs) ^9–11^. ISCs divide to produce an enteroblast (EB) about 90-95% of the time ^11–13^. The EB receives a high Notch signal from the ISC and differentiates into an EC ^12, 14^. On the other hand, EEs arise from ISCs via an enteroendocrine precursor (EEp) and by direct ISC differentiation into an EE ^15^. Together, ISCs, EBs and EEps are considered adult midgut progenitors and express the progenitor marker, *escargot* (*esg*) ^10^. The fly midgut epithelium is surrounded by two layers of visceral muscle (VM) and oxygen-carrying trachea ^11, 16, 17^. Together, the EBs, ECs, EEs, the VM and trachea act as a regenerative niche after midgut damage and produce cues that directly stimulate ISC proliferation. These cues include the Unpaired cytokines Upd1-3, EGFs, Wnt, Hedgehog, Fibroblast Growth Factor and PTK7/Off-track ^16–25^.

Many of the signals received by fly ISCs are also received by mammalian ISCs from the regenerative niche after damage. One such niche-derived cue promoting regeneration of both the *Drosophila* and mammalian intestines is cytokine-Stat signalling. STAT (Stat92E) activity in *Drosophila* ISCs is required for their proliferation after damage caused by pathogenic bacteria ^18, 26^. STAT activation within progenitors is thought to be exclusively through cytokine-like Unpaireds (Upd1-3) produced by enteroblasts and damaged enterocytes upon pathogenic bacterial infection ^19–21, 23^. In the mammalian intestine, Stat3 is activated after epithelial damage by either γ-irradiation or infection by the mouse pathogen, *Citrobacter* rodentium ^27, 28^ and required for epithelial regeneration after intestinal damage ^29, 30^. Stat3 is thought to be partly activated by cytokines produced by macrophages and type 3 innate lymphoid cells ^5, 6^.

Whilst much is known about the signals received by ISCs from the regenerative niche upon damage, some recent evidence from both invertebrates and vertebrates has shown that epithelial damage sensing directly by stem cells themselves is essential for stem cell-mediated regeneration ^31, 32^. In the wounded mouse skin, epidermal stem cells at the wound margin detect hypoxia caused by local blood vessel damage and respond by producing the cytokine, IL-24. This cytokine activates Stat3 in epidermal stem cells, stimulating their proliferation ^31^. Similarly, in the adult *Drosophila* and mouse intestines, ISCs sense an increase in very long-chain fatty acids within the niche after injury. This is mediated by peroxisome proliferator-activated receptors (PPARs), which promote differentiation by increasing peroxisome numbers and Sox21 activity ^32^. These studies strongly indicate that stem cells can directly sense tissue damage and activate intrinsic mechanisms that promote regeneration without cues from the regenerative niche.

It remains unclear how, mechanistically, ISCs sense intestinal damage and translate this into their proliferation to regenerate the intestinal epithelium. The non-receptor tyrosine kinase Src is a crucial intrinsic factor required by both adult *Drosophila* and mammalian ISCs in response to tissue damage to induce ISC proliferation. In adult *Drosophila*, Src42A signalling is required in progenitors for ISC proliferation after pathogenic bacterial infection ^27, 33^. The *Drosophila* genome encodes two Src family kinases, Src42A and Src64B. Whereas Src64B expression can stimulate ISC proliferation in unchallenged midguts, its requirement for ISC proliferation after midgut damage is not clear ^27, 33^. Src has been shown to be activated in progenitors by Wnt/Wingless (Wg) signalling after pathogenic bacterial infection ^27^ and by the overexpression of the receptor tyrosine kinase, Ret ^34^. Constitutive activation of Src42A in progenitors strongly induces ISC proliferation via EGFR/ERK signalling and STAT ^27, 33^. JAK-STAT signalling is usually activated by Unpaired (Upd1-3) cytokines that bind to the Domeless receptor ^35^. Interestingly, Domeless and its ligands, Upd1-3, are not required for Src42A to stimulate ISC proliferation through STAT activation ^27, 33^. Therefore, how STAT is activated by Src42A in midgut progenitors is not understood. Similarly in the mouse intestine, both Src and Stat3 are activated after γ-irradiation, and Stat3 activation after intestinal damage partly required Src ^27^. However, it is not entirely understood in both the mouse intestine and the adult *Drosophila* midgut how Src is activated after damage and how Src activates Stat3 and Stat92E, respectively.

In *Drosophila*, Src signalling can activate STAT via Shark signalling. To do so, Src kinases phosphorylate tyrosines within immunoreceptor tyrosine-based activation motifs (ITAMs) on immune receptors, such as Draper ^36^. Phosphorylation of tyrosine residues within ITAMs then allows the non-receptor tyrosine kinase, Spleen tyrosine kinase (Syk)—Shark in *Drosophila*—to dock onto ITAMs in Draper via their Src-homology 2 (SH2)-domains. This allows for full Syk activation by enabling phosphorylation of the linker region between its SH2 and kinase domains by Src family kinases ^36, 37^. In *Drosophila,* phagocytic macrophage-like hemocytes use Draper to detect and engulf apoptotic corpses ^38, 39^, and Draper-Src-Shark signalling and Draper-mediated engulfment are also used by glial cells to clear severed axons from neurons after injury ^38, 40, 41^. Furthermore, glia activate STAT through Draper-Src-Shark signalling after adult *Drosophila* olfactory receptor neuron (ORN) injury ^42^. Draper recognises dying cells through binding of exposed phosphatidylserine on the outside surface of apoptotic cells, which promotes Draper phosphorylation and engulfment signalling ^43, 44^.

Thus, Draper-Src-Shark signalling might be utilised by ISCs to detect apoptotic cells within damaged midguts and couple this to ISC proliferation, and such a role of Draper-Src-Shark-STAT signalling has not yet been explored. While the mammalian Draper homologue, MEGF10, is required in muscle satellite cells for skeletal muscle regeneration ^45^, very little is known about the role of Syk or MEGF10 in stem cells during regeneration.

Here we explore how adult *Drosophila* midgut ISCs sense damage after midgut injury and translate this into a regenerative response. We find that Draper-Src-Shark signalling is required within ISCs for adult *Drosophila* midgut regeneration by recognising midgut damage and translating it to ISC proliferation. Mechanistically, Draper-Src-Shark signalling stimulates ISC proliferation by increasing STAT activity in ISCs, which has previously been thought to be exclusively activated by cytokine (Upd1-3)/JAK signalling. Furthermore, we show that Draper-Src-Shark signalling in progenitors activates STAT in the visceral muscle, demonstrating how progenitors shape the regenerative niche to support epithelial regeneration.

## Results

### Src induces proliferation in adult *Drosophila* intestinal stem cells after midgut damage

Oral infection of adult flies with gram-negative pathogenic bacteria (*Pseudomonas entomophila*, *P.e.* or *Erwinia caratovora caratovora 15*, *Ecc15*) has been shown to increase the levels of activated Src in the midgut epithelium, including within *esg^+^* progenitors ^27, 46^. However, the subcellular localisation of activated Src was not clear. Using an antibody recognising activated Src (phosphorylated tyrosine 419 on human Src, pSrc)^34^, we found that Src is activated after *P.e.* infection in *esg*^+^ progenitors and in ECs (Fig. 1A-C; Supplementary Fig. 1D-E’). Moreover, we found a strong cortical increase by 74.2% of activated Src in *esg^+^* progenitors (Fig. 1C), which partially overlapped with Armadillo (Arm). High levels of Arm are found at the cortex of adult midgut progenitors (Fig. 1A, B)^47^. We validated that the pSrc antibody is specific for activated Src42A: Whereas depletion of Src42A in ECs decreased pSrc levels in ECs after *P.e.* infection, activated Src42A levels still increased in progenitors (Supplementary Fig. 1D-G’). These data suggest that activated Src42A accumulates at the cortex of progenitors after pathogenic bacterial infection.

**Figure 1.**
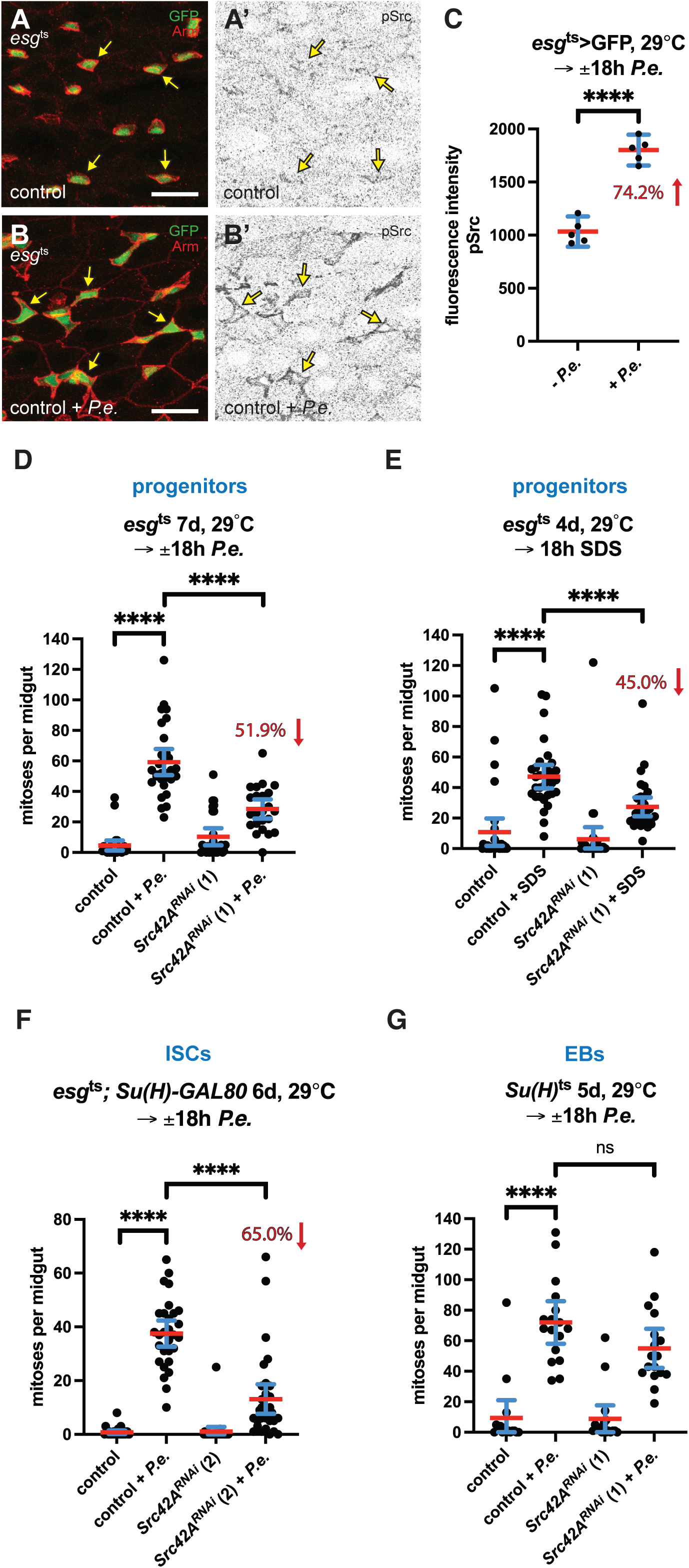
Src signalling is required in adult *Drosophila* intestinal stem cells for their proliferation after midgut damage. Activated Src accumulates cortically in progenitors after *P.e.* infection **(A-C)**. Phosphorylated Src Y419 (pSrc) levels increased in progenitors (*esg*>GFP^+^; green, Arm^high+^; red, yellow arrows) of *P.e.*-infected midguts (**B, B’**) compared to those in uninfected midguts (**A, A’**). The mean cortical pSrc levels increased in progenitors of *P.e.*-infected midguts (n=5 midguts from one of two experiments) compared to uninfected control midguts (n=5 midguts from one of two experiments) (unpaired *t* test; *P* < 0.0001) (**C**). Src42A is required in progenitors to promote intestinal stem cell (ISC) proliferation after *P.e.* infection **(D)**. The number of mitoses increased in *P.e.*-infected control midguts (n=29 midguts from two experiments) compared to uninfected control midguts (n=27 midguts from two experiments) (Mann-Whitney; *P* < 0.0001). ISC proliferation was suppressed by 51.9% in *P.e.*-infected midguts expressing *Src42A^RNAi^* in progenitors for 7 days with *esg*^ts^ (n=22 midguts from two experiments) compared to *P.e.*-infected control midguts (Mann-Whitney; *P* < 0.0001). Src42A is required in progenitors to promote ISC proliferation after SDS-induced damage **(E)**. The number of mitoses increased in SDS-fed control midguts (n=30 midguts from two experiments) compared to undamaged control midguts (n=30 midguts from two experiments) (Mann-Whitney; *P* < 0.0001). ISC proliferation was suppressed by 45.0% in SDS-fed midguts expressing *Src42A^RNAi^* in progenitors for 4 days with *esg*^ts^ (n=30 midguts from two experiments) compared to SDS-fed control midguts (Mann-Whitney; *P* < 0.0001). Src42A is required in ISCs to promote their proliferation after *P.e.* infection **(F)**. The number of mitoses increased in *P.e.*-infected control midguts (n=29 midguts from two experiments) compared to uninfected control midguts (n=29 midguts from two experiments) (Mann-Whitney; *P* < 0.0001). ISC proliferation was suppressed by 65.0% in *P.e.*-infected midguts expressing *Src42A^RNAi^* in ISCs for 6 days with *esg*^ts^ *Su(H)-GAL80* (n=32 midguts from two experiments) compared to *P.e.*-infected control midguts (Mann-Whitney; *P* < 0.0001). Src42A is not required in enteroblasts (EBs) to promote ISC proliferation after *P.e.* infection **(G)**. The number of mitoses increased in *P.e.*-infected control midguts (n=17 midguts from one experiment) compared to uninfected control midguts (n=16 midguts from one experiment) (Mann-Whitney; *P* < 0.0001). ISC proliferation was not significantly blocked in *P.e.*-infected midguts expressing *Src42A^RNAi^* in EBs for 5 days with *Su(H)*^ts^ (n=17 midguts from one experiment) compared to *P.e.*- infected control midguts (Mann-Whitney; *P* < 0.0585). Scale bars in A-B are 20μm.

Src42A is required in midgut progenitors for ISC proliferation after *P.e.* infection ^27, 33^; however, it is not known whether Src42A senses midgut damage or pathogenic bacteria. We confirmed that depletion of Src42A from progenitors by RNAi with the progenitor-specific driver *esg-GAL4 tubGAL80*^ts^ (*esg*^ts^) blocked ISC proliferation after *P.e.* infection by 51.9% and 77.8%, for RNAis (1) and (2), respectively (Fig 1D, Supplementary Fig. 1B). This decrease in ISC proliferation was not caused by the loss of progenitors as the mean percent *esg-*positive progenitors in these midguts was similar to uninfected and *P.e*-infected control midguts (Supplementary Fig. 1A). We next determined whether Src42A detected pathogenic bacteria (*P.e.*) or rather tissue damage from infection. To ascertain this, we depleted Src42A in progenitors with *esg*^ts^ and fed these flies sodium dodecyl sulfate (SDS). While SDS exposure increased ISC proliferation in control midguts, we found that Src42A depletion in progenitors blocked ISC proliferation by 45.0% after detergent-induced damage (Fig. 1E). Together these data suggest that Src42A is required in progenitors to sense midgut damage rather than pathogenic bacteria themselves.

*esg^+^* midgut progenitors include ISCs, EBs and EEps. Thus, it was uncertain in which progenitor cell type Src42A was required for ISC proliferation after *P.e.*-induced damage. To determine this, we depleted Src42A from ISCs using an ISC-specific driver, *esgGAL4 Su(H)Gal80 tubGAL80*^ts^ (*esg*^ts^ *Su(H)GAL80*) and in EBs using an EB-specific driver, *Su(H)GAL4 tubGal80*^ts^. We found that Src42A depletion in ISCs strongly blocked ISC proliferation after *P.e.* by 65.0% (Fig. 1F). In contrast, we found that EB-specific depletion of Src42A did not significantly decrease ISC proliferation after *P.e.* infection (Fig. 1G). These data suggest that Src42A acts mainly in ISCs to respond to midgut damage.

### Draper-Src-Shark signalling promotes ISC proliferation after damage

The expression of constitutively active Src42A in midgut progenitors strongly induces ISC proliferation through EGFR/ERK signalling and STAT ^27, 33^. However, the mechanism by which Src42A regulates STAT is unknown. In adult *Drosophila* glia, Src42A signalling can activate STAT via Draper-Shark signalling to promote phagocytosis of injured axons of ORNs ^42^. Thus, we tested whether Draper and Shark are required for ISC proliferation after damage by depleting both *draper* and *shark* from midgut progenitors with *esg*^ts^. Similar to Src42A depletion, we found that both Draper and Shark are required in progenitors for ISC proliferation after *P.e.* infection. Depleting *shark* in progenitors by RNAi with *esg*^ts^ resulted in a 51.0% and 52.4% (RNAi (1) and (2), respectively) decrease in ISC proliferation after *P.e.* infection compared to infected control midguts (Fig. 3A, Supplementary Fig. 2B). Furthermore, depleting *draper* from progenitors with *esg*^ts^ resulted in a 48.4% and 44.7% (RNAi (1) and RNAi (2), respectively) decrease in ISC proliferation after *P.e.* infection compared to infected control midguts (Fig. 4A, Supplementary Fig. 3B). Moreover, the decrease in ISC proliferation after depleting Draper and Shark within progenitors and *P.e.* infection was not due to reduced progenitor cell number (Supplementary Fig. 2A, 3A). Together these data indicate that Draper-Src-Shark signalling in progenitors promotes adult *Drosophila* midgut regeneration upon damage.

**Figure 2.**
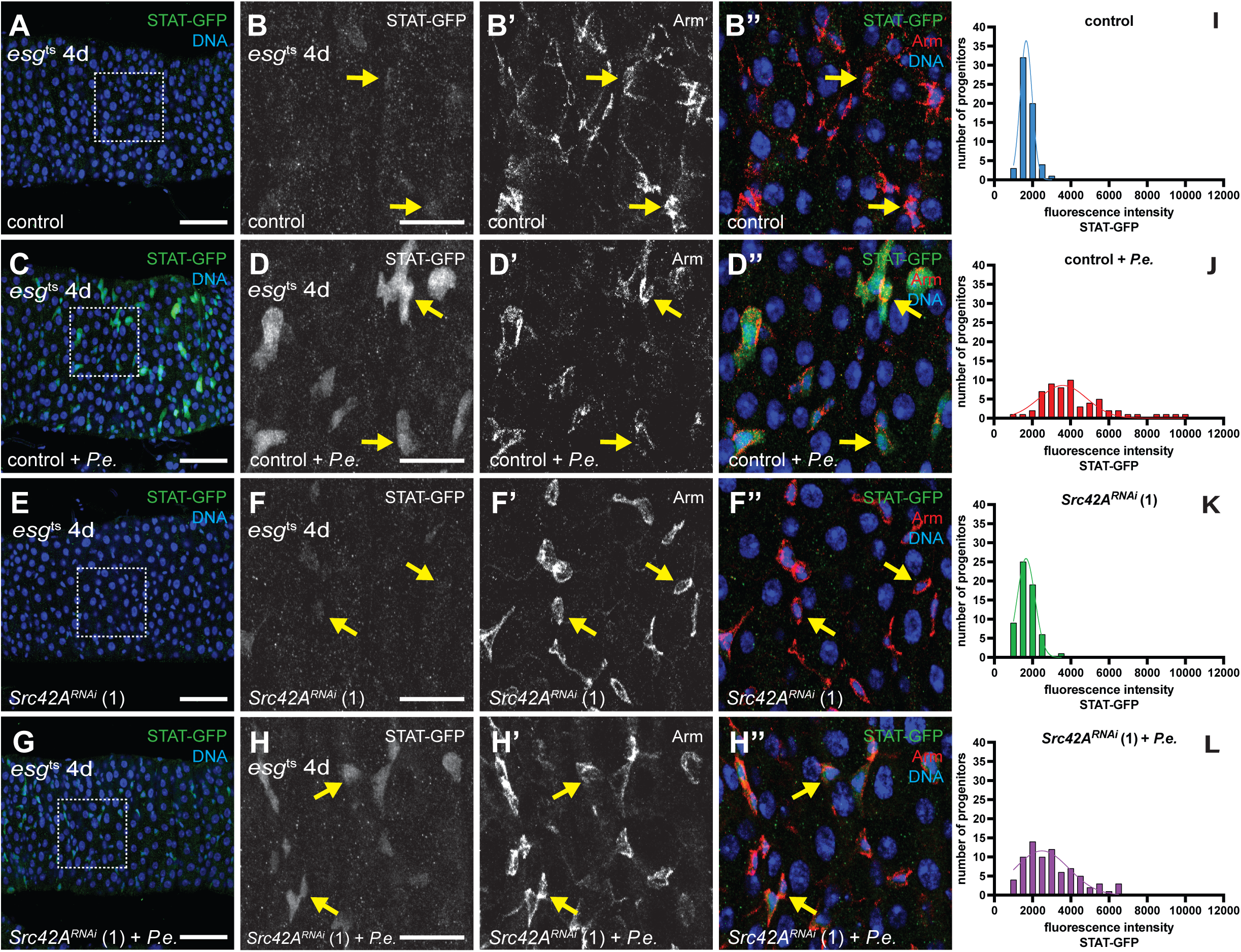
Src signalling stimulates Stat activity in progenitors after pathogenic bacteria-induced damage. Stat is activated in progenitors (*esg*^+^ Arm^high+^; red; yellow arrows) after *P.e.* infection **(A-D”, I-J)**. Stat activity (10XSTAT-GFP; green) increased in progenitors of *P.e.*-infected control midguts (**C, D-D’’**) compared to uninfected control midguts (**A, B-B’’**). Stat activity (STAT-GFP fluorescence intensity) increased in progenitors of *P.e.*-infected control midguts (n=60 cells from four midguts from one of two experiments) (**J**) compared to uninfected control midguts (n=60 cells from four midguts from one of two experiments) (**I**). Src42A is required to activate Stat in progenitors after *P.e.* infection **(E-H”, K-L)**. Stat activation decreased in progenitors of *P.e.*-infected midguts expressing *Src42A^RNAi^* with *esg*^ts^ (**G, H-H’’**) compared to *P.e.*- infected control midguts (**C, D-D’’**). Stat activity decreased in progenitors from *P.e.*- infected midguts expressing *Src42A^RNAi^* in progenitors for 4 days using *esg*^ts^ (n=60 cells from four midguts from one of two experiments) (**L**) compared to infected control midguts (**J**). Stat activation did not significantly change in uninfected midguts expressing *Src42A^RNAi^* for 4 days with *esg*^ts^ (n=60 cells from four midguts from one of two experiments) (**K**) compared to uninfected control midguts (**I**). DNA is in blue in A, C, E, G and B’’, D’’, F’’ and H’’. Scale bars in A, C, E and G are 50μm and in B, D, F and H are 20μm.

**Figure 3.**
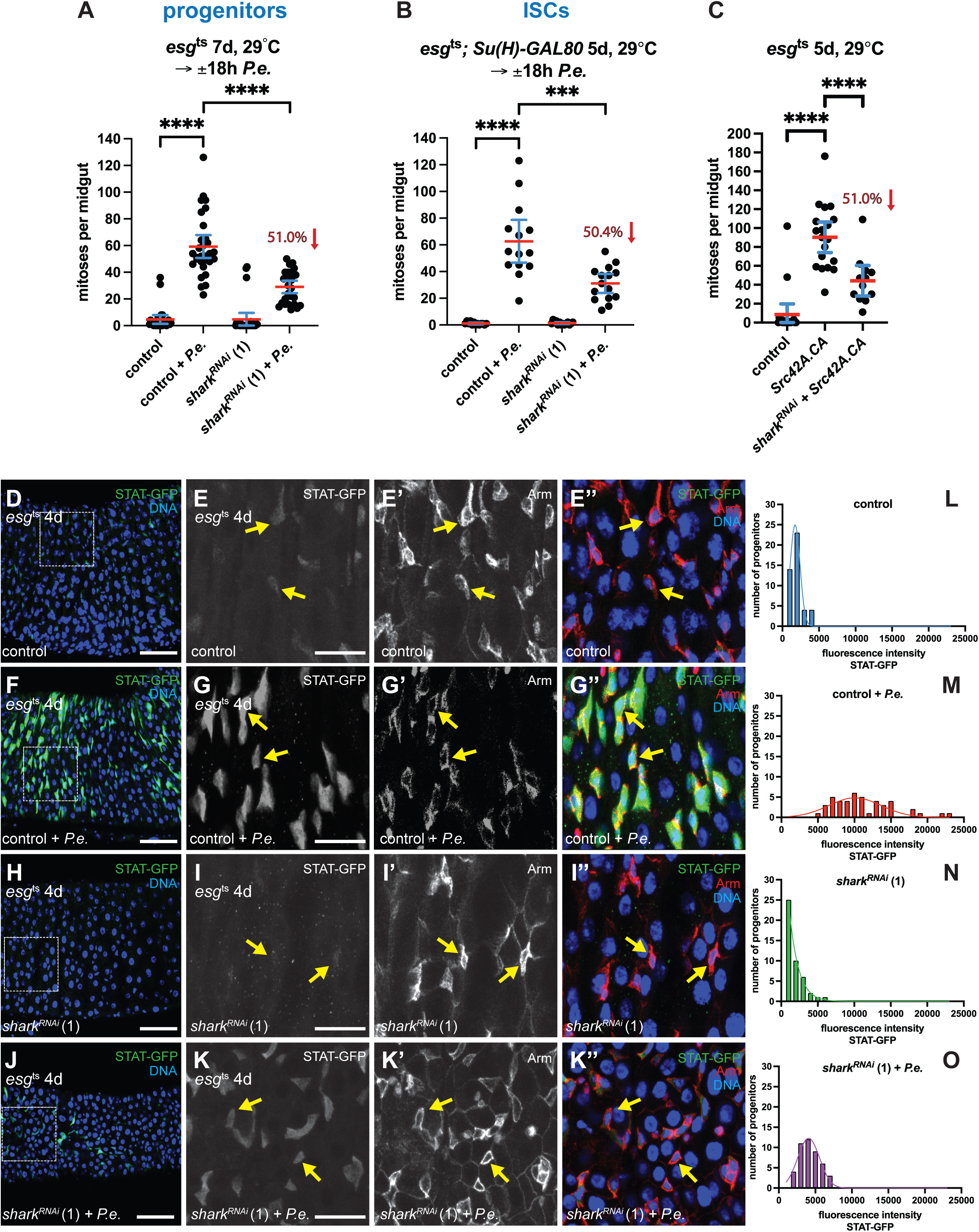
The Syk-homologue, Shark, is required for intestinal stem cell proliferation and for activating Stat in progenitors following pathogenic bacteria-induced damage. Shark is required in progenitors to promote ISC proliferation after *P.e.* infection **(A)**. The number of mitoses increased in *P.e.*-infected control midguts (n=29 midguts from two experiments) compared to uninfected control midguts (n=27 midguts from two experiments) (Mann-Whitney; *P* < 0.0001). ISC proliferation was suppressed by 51.0% in *P.e.*-infected midguts expressing *shark^RNAi^* in progenitors for 7 days with *esg*^ts^ (n=27 midguts from two experiments), compared to *P.e.*-infected control midguts (Mann-Whitney; *P* < 0.0001). Shark is specifically required in ISCs to promote their proliferation after *P.e.* infection **(B)**. The number of mitoses increased in *P.e.*-infected control midguts (n=14 midguts from one of two experiments) compared to uninfected control midguts (n=13 midguts from one of two experiments) (Mann-Whitney; *P* < 0.0001). ISC proliferation was suppressed by 50.4% in *P.e.*-infected midguts expressing *shark^RNAi^* in ISCs for 5 days with *esg*^ts^ *Su(H)-GAL80* (n=15 midguts from one of two experiments), compared to *P.e.*-infected control midguts (Mann-Whitney; *P* = 0.0003). Shark is required for Src42A to promote ISC proliferation **(C)**. The number of mitoses increased in midguts expressing constitutive active Src42A (*Src42A.CA*) (n=19 midguts from one of two experiments) compared to control midguts (n=20 midguts from one of two experiments) (Mann-Whitney; *P* < 0.0001). ISC proliferation was suppressed by 51.0% in midguts expressing *shark^RNAi^* together with *Src42A.CA* (n=12 midguts from one of two experiments) compared to *Src42A.CA*-expressing midguts (Mann-Whitney; *P* < 0.0001). Stat is activated in progenitors (*esg*^+^ Arm^high+^; red; yellow arrows) after *P.e.* infection **(D-G”, L-M)**. Stat activity (STAT-GFP; green) increased in progenitors of *P.e.*- infected control midguts (**F, G-G’’**) compared to uninfected control midguts (**D, E-E’’**). Stat activity (10X STAT-GFP fluorescence intensity) increased in progenitors of *P.e.*- infected control midguts (n=45 cells from three midguts from one of three experiments) (**M**) compared to uninfected control midguts (n=45 cells from three midguts from one of three experiments) (**L**). Shark is required to activate Stat in progenitors after *P.e.* infection **(H-K”, N-O)**. Stat activation decreased in progenitors of *P.e.*-infected midguts expressing *shark^RNAi^* in progenitors for 4 days using *esg*^ts^ (**J, K-K’’**) compared to *P.e.*- infected control midguts (**F, G-G’’**). Stat activity decreased in progenitors from *P.e.*- infected midguts expressing *shark^RNAi^* in progenitors for 4 days with *esg*^ts^ (n=45 cells from three midguts from one of three experiments) (**O**) compared to infected control midguts (**M**). Stat activation was not significantly blocked in uninfected midguts expressing *shark^RNAi^* for 4 days with *esg*^ts^ (n=45 cells from three midguts from one of three experiments) (**N**) compared to uninfected control midguts (**L**). DNA is in blue in D, F, H and J and E’’, G’’, I’’ and K’’. Scale bars in D, F, H and J are 50μm and in E, G, I and K are 20μm.

**Figure 4.**
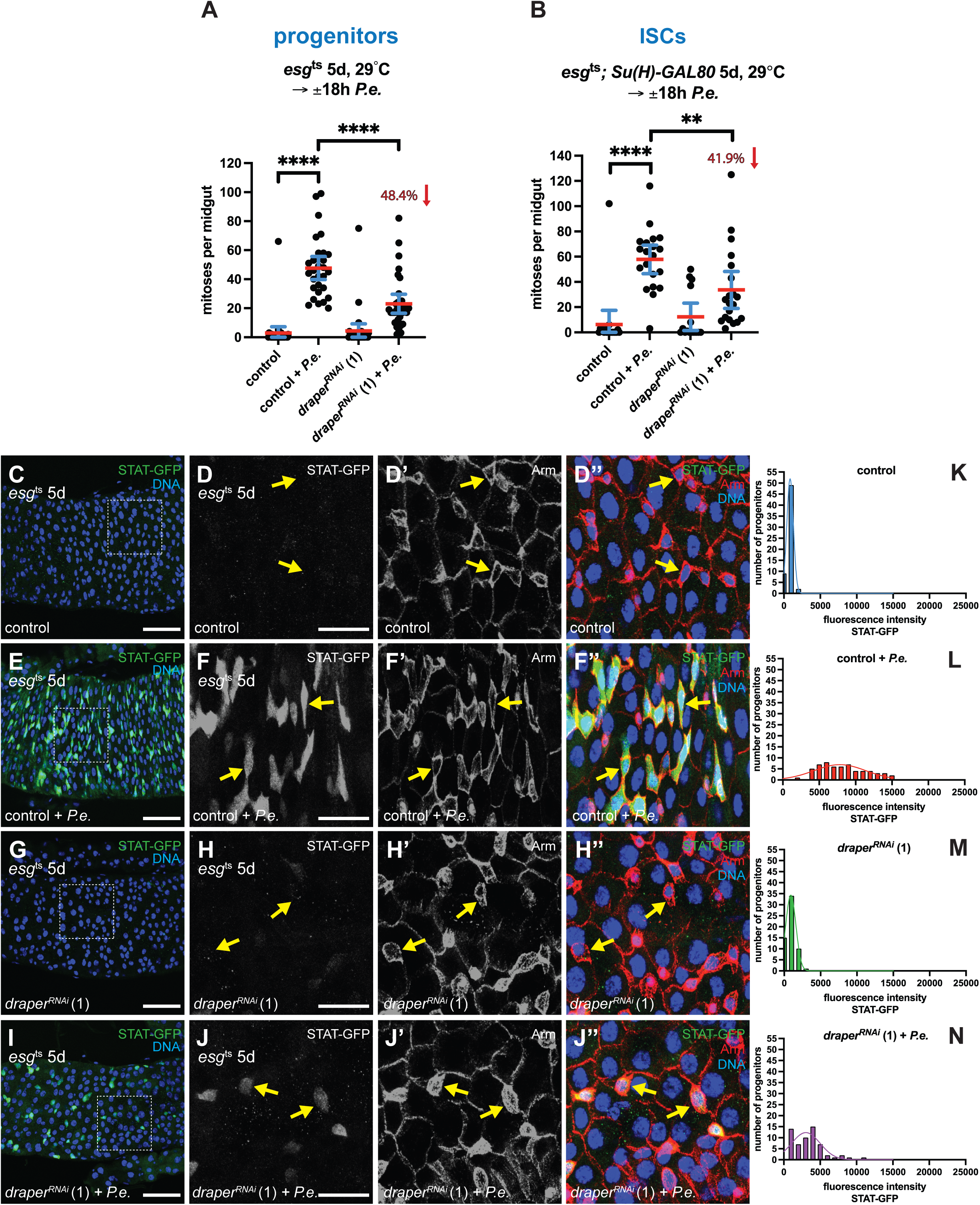
Draper is required for intestinal stem cell proliferation and for activating Stat in progenitors after pathogenic bacteria-induced damage. Draper is required in progenitors to promote ISC proliferation after *P.e.* infection **(A)**. The number of mitoses increased in *P.e.*-infected control midguts (n=29 midguts from two experiments) compared to uninfected control midguts (n=31 midguts from two experiments) (Mann-Whitney; *P* < 0.0001). ISC proliferation was suppressed by 48.4% in *P.e.*-infected midguts expressing *draper^RNAi^* in progenitors for 5 days with *esg*^ts^ (n=32 midguts from two experiments) compared to *P.e.*-infected control midguts (Mann-Whitney; *P* < 0.0001). Draper is specifically required in ISCs to promote their proliferation after *P.e.* infection **(B)**. The number of mitoses increased in *P.e.*-infected control midguts (n=20 midguts from one of two experiments) compared to uninfected control midguts (n=19 midguts from one of two experiments) (Mann-Whitney; *P* < 0.0001). ISC proliferation was suppressed by 41.9% in *P.e.*-infected midguts expressing *draper^RNAi^* in ISCs for 5 days with *esg*^ts^ *Su(H)-GAL80* (n=20 midguts from one of two experiments) compared to *P.e.*- infected control midguts (Mann-Whitney; *P* = 0.0026). Stat is activated in progenitors (*esg*^+^, Arm^high+^; red; yellow arrows) after *P.e.* infection **(C-F”, K-L)**. Stat activity (STAT-GFP; green) increased in progenitors of *P.e.*-infected control midguts (**E, F-F’’**) compared to uninfected control midguts (**C, D-D’’**). Stat activity (10X STAT-GFP fluorescence intensity) increased in progenitors of *P.e.*-infected control midguts (n=60 cells from four midguts from one of two experiments) (**L**) compared to uninfected control midguts (n=60 cells from four midguts from one of two experiments) (**K**). Draper is required to activate Stat in progenitors after *P.e.* infection **(G-J”, M-N)**. Stat activation decreased in progenitors of *P.e.*-infected midguts expressing *draper^RNAi^* in progenitors for 5 days with *esg*^ts^ (**I, J-J’’**) compared to *P.e.*-infected control midguts (**E**, **F**-**F’’**). Stat activity decreased in progenitors from *P.e.*-infected midguts expressing *draper^RNAi^* in progenitors for 5 days with *esg*^ts^ (n=60 cells from four midguts from one of two experiments) (**N**) compared to infected control midguts (**L**). Stat activation was not significantly blocked in uninfected midguts expressing *draper^RNAi^* for 5 days with *esg*^ts^ (n=60 cells from four midguts from one of two experiments) (**M**) compared to uninfected control midguts (**K**). DNA is in blue in C, E, G and I and D’’, F’’, H’’ and J’’. Scale bars in C, E, G and I are 50μm and in D, F, H and J are 20μm.

To determine whether Shark and Draper are required in ISCs to induce ISC proliferation after midgut damage, we depleted Shark and Draper from ISCs using *esg*^ts^ *Su(H)Gal80*. We found that Shark and Draper depletion in ISCs strongly blocked ISC proliferation after *P.e.* by 50.4% and 41.9%, respectively (Fig. 3B, 4B). These data indicate that similar to Src42A, Shark and Draper act mainly in ISCs to respond to midgut damage.

Src family kinases are inhibited by C-terminal tyrosine phosphorylation by the C-terminal Src kinase (Csk). Expression of a constitutively active Src42A (Y511F), which is refractory to Csk, in progenitors with *esg*^ts^ stimulates ISC proliferation in the absence of damage ^27, 33^. To determine whether Shark is required for Src42A to promote ISC proliferation, we depleted *shark* in progenitors with RNAi and simultaneously expressed constitutively active Src42A in progenitors using *esg*^ts^. As expected, expression of constitutively active Src42A in progenitors stimulated ISC proliferation (Fig. 3C). However, in combination with an RNAi towards *shark* in progenitors, ISC proliferation was blocked by 51.0% compared to constitutively active Src42A expression alone (Fig. 3C). These data suggest that Draper-Shark signalling is partially required for Src42A to promote ISC proliferation.

### Draper-Src-Shark signalling promotes ISC proliferation through STAT activation

JAK-STAT signalling is crucial for adult *Drosophila* ISCs to proliferate after midgut damage by pathogenic bacteria ^18, 26^. STAT activity can be observed in the midgut using the reporter, *10XSTAT-DGFP* (Stat-GFP)^18, 48, 49^. This reporter consists of multimerized 10X STAT92E binding sites fused together with a destabilized GFP. STAT activity has been observed in progenitors in homeostatic midguts but not within ECs and EEs ^18, 48^. Further, JAK-STAT signalling increases within midgut progenitors and the visceral muscle after pathogenic bacterial infection ^18, 20, 21^.

Given that expression of constitutively active Src42A in progenitors induces ISC proliferation, which requires STAT ^27, 33^, we wondered whether STAT activity is directly regulated by Src42A. To test whether Src42A, Shark and Draper are required for STAT activation in progenitors after *P.e.* infection, we depleted *Src42A*, *shark* or *draper* from progenitors with *esg*^ts^ and infected flies with *P.e.*. Midgut progenitors can be identified by their high levels of cortical ß-catenin/Armadillo (Arm), whereas Arm is found at the adherens junctions of EEs and ECs ^47, 50^. We therefore measured STAT-GFP within *esg^+^* Arm^high+^ progenitors. We found an increase in STAT activity in the posterior midgut (R4-R5) after *P.e.* infection. Whereas Stat activity increased in the progenitors in *P.e.*-infected control midguts (Fig. 2C,D-D’’, J; Fig. 3F, G-G’’, M; Fig. 4E, F-F’’, L; Supplementary Fig. 1C, 2C, 3C) compared to uninfected control midguts (Fig. 2A, B-B’’, I; Fig. 3D, E-E”, L; Fig. 4C, D-D’’, K; Supplementary Fig. 1C, 2C, 3C), we found that Stat activity did not similarly increase in *Src42A-*, *shark*- and *draper*-depleted progenitors in *P.e.-*infected midguts (Fig. 2G, H-H’’, L; Fig. 3J, K-K”, O; Fig. 4I, J-J’’, N; Supplementary Fig. 1C, 2C, 3C). Indeed, we found a 41.4%, 62.6% and 58.7% decrease in STAT activity in *Src42A*-, *shark*- and *draper*-depleted progenitors, respectively, after *P.e.* infection (Supplementary Fig. 1C, 2C, 3C). These data indicate that Draper-Src-Shark signalling promotes ISC proliferation through STAT activation. Interestingly, STAT activation decreased in both *esg^+^* Arm^high+^ ISCs and EBs. These data further suggest that Draper-Src-Shark signalling regulates STAT activity in EBs either cell-autonomously, or in a cell non-autonomous manner via signalling from ISCs to EBs. Furthermore, we found similar levels of STAT activity in *Src42A*-, *shark*- and *draper*-depleted progenitors of uninfected midguts (Fig. 2E, F-F’’, K; Fig. 3H, I-I”, N; Fig. 4G, H-H’’, M; Supplementary Figure 1C, 2C, 3C) as to progenitors in uninfected control midguts. These data suggest that Draper-Src-Shark signalling is not required for STAT activity and ISC proliferation during homeostasis and rather functions exclusively as a damage-sensing pathway. We next examined STAT activation in the visceral muscle (VM)—part of the regenerative ISC niche—of infected midguts. Compared to control uninfected midguts, STAT activity increased within midgut VM upon *P.e.*-infection (Fig. 6A-B’, 5E-F’, I-K; Supplementary Fig. 1H-I’). Surprisingly, we found a 37.1%, 57.9% and 27.0% decrease in STAT activity in the VM of *P.e.*-infected midguts containing *Src42A-*, *shark*- and *draper*-depleted progenitors, respectively, compared to *P.e.*-infected control midguts (Fig. 6B-B’, D-D’, F-F’, H-H’, I-K; Supplementary Fig. 1I-I’, K-K’). These data indicate that STAT activity in the VM is regulated cell non-autonomously by Draper-Src-Shark signalling in ISCs, EBs, or both ISCs and EBs after midgut damage.

**Figure 5.**
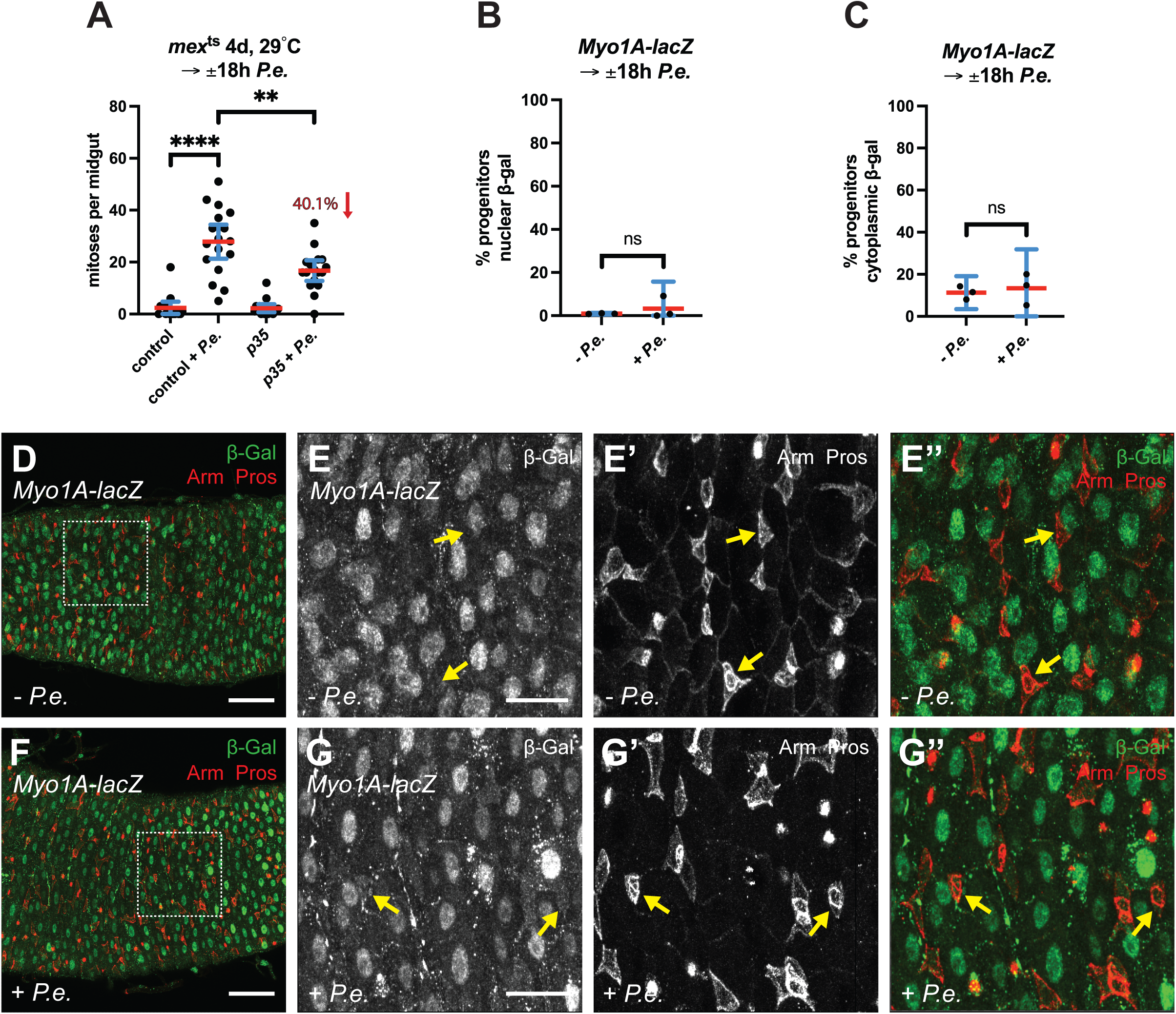
Apoptotic enterocytes promote intestinal stem cell proliferation without their engulfment following pathogenic bacteria-induced damage. Blocking apoptosis partially inhibits ISC proliferation after *P.e.* infection **(A)**. The number of mitoses increased in *P.e.*-infected control midguts (n=17 midguts from one experiment) compared to uninfected control midguts (n=16 midguts from one experiment) (Mann-Whitney; *P* < 0.0001). ISC proliferation decreased by 40.1% in *P.e.*-infected midguts overexpressing p35 in enterocytes (ECs) for 4 days with *mex*^ts^ (n=17 midguts from one experiment) compared to *P.e.*-infected control midguts (Mann-Whitney; *P* = 0.0052). Progenitors did not engulf apoptotic ECs after *P.e.* infection **(B-G’’)**. Nuclear localised β-gal expression (green) was not induced within Arm^high+^ (red) progenitors after *P.e.* infection as the mean percent of total progenitors with nuclear β-gal did not significantly differ between control and *P.e.*-infected midguts (**B; D, E-E’’; F, G-G’’;** yellow arrows). β-Gal, possibly from neighbouring ECs, was observed within the cytoplasm of Arm^high+^ progenitors at the same frequency in both control and *P.e.*- infected midguts (**C; D, E-E’’; F, G-G’’**). Scale bars in D and F are 50μm and in E and G are 20μm.

**Figure 6.**
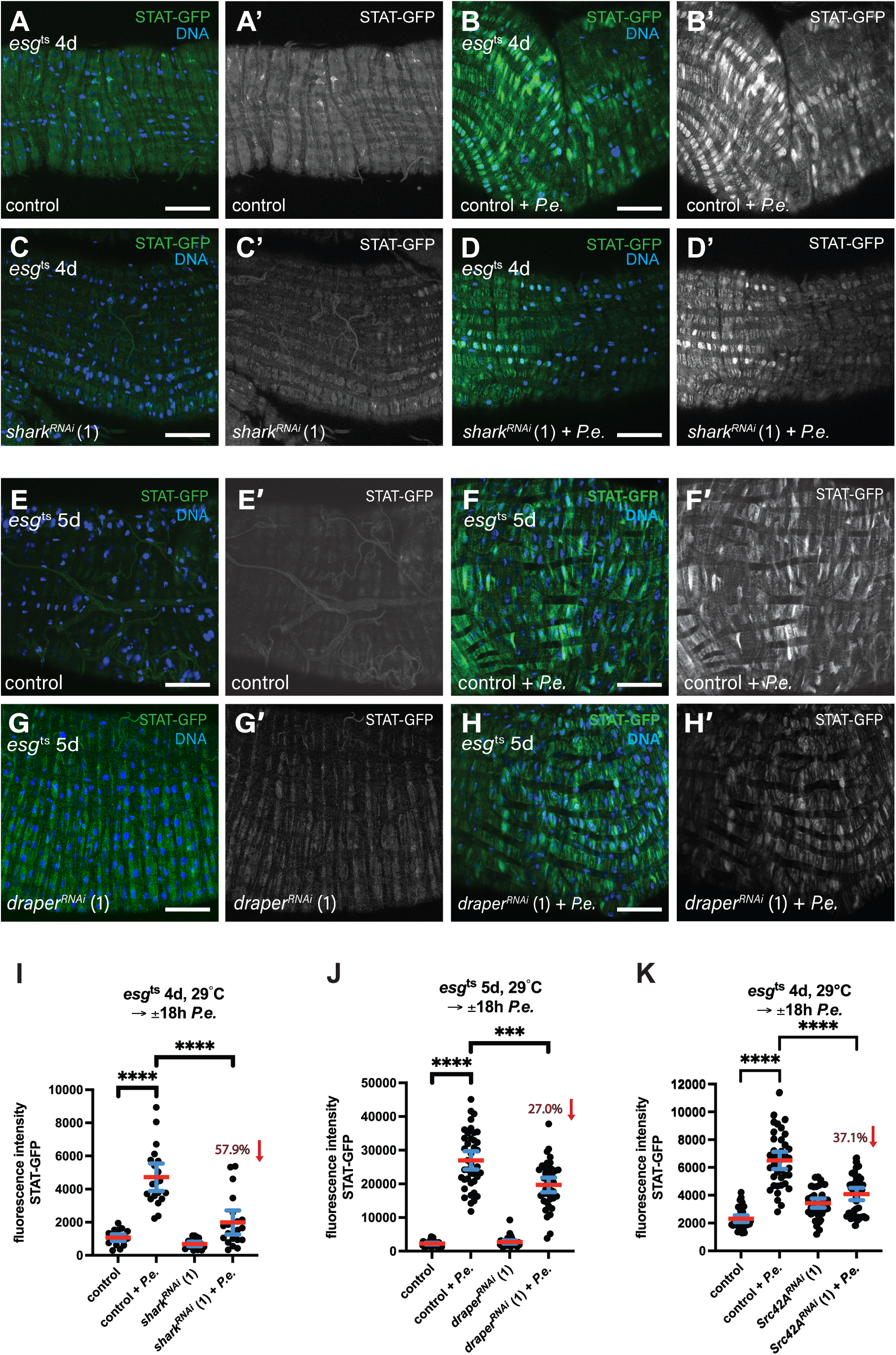
Draper-Src-Shark signalling in progenitors activates Stat in the visceral muscle after pathogenic bacteria-induced damage. Shark is required to activate Stat in the visceral muscle (VM) after *P.e.* infection **(A-D, I)**. Stat activity (green) increased in the VM of *P.e.*-infected control midguts (**B-B’**) compared to uninfected control midguts (**A-A’**). Stat activation decreased in the VM of *P.e.*-infected midguts expressing *shark^RNAi^* for 4 days with *esg*^ts^ (**D-D’**) compared to *P.e.*-infected control midguts. Mean Stat activity (10X STAT-GFP fluorescence intensity) increased in the VM of *P.e.*-infected control midguts (n=20 cells from two midguts from one of three experiments) compared to uninfected control midguts (n=20 cells from two midguts from one of three experiments) (Mann-Whitney; *P* < 0.0001) (**I**). Mean Stat levels decreased in the VM of *P.e.*-infected midguts expressing *shark^RNAi^* in progenitors with *esg*^ts^ by 57.9% (n=20 cells from two midguts from one of three experiments) compared to infected control midguts (Mann-Whitney; *P* < 0.0001) (**I**). Draper is required to activate Stat in the VM after *P.e.* infection **(E-H’, J)**. Stat activity increased in the VM of *P.e.*-infected control midguts (**F-F’**) compared to uninfected control midguts (**E-E’**). Stat activation decreased in the VM of *P.e.*-infected midguts expressing *draper^RNAi^* for 5 days with *esg*^ts^ (**H-H’**) compared to *P.e.*- infected control midguts. Mean Stat activity (10X STAT-GFP fluorescence intensity) increased in the VM of *P.e.*-infected control midguts (n=40 cells from four midguts from one experiment) compared to uninfected control midguts (n=40 cells from four midguts from one experiment) (Mann-Whitney; *P* < 0.0001) (**J**). Mean Stat levels decreased in the VM of *P.e.*-infected midguts expressing *draper^RNAi^* in progenitors with *esg*^ts^ by 27.0% (n=40 cells from four midguts from one experiment) compared to infected control midguts (Mann-Whitney; *P* = 0.0003) (**J**). Src42A is required to activate Stat in the VM after *P.e.* infection **(K)**. Mean Stat activity increased in the VM of *P.e.*-infected control midguts (n=40 cells from four midguts from one of two experiments) compared to uninfected control midguts (n=40 cells from four midguts from one of two experiments) (Mann-Whitney; *P*<0.0001). Mean Stat activity decreased in the VM of *P.e.*-infected midguts expressing *Src42A^RNAi^* by 37.1% (n=40 cells from four midguts from one of two experiments) compared to *P.e.*-infected control midguts (Mann-Whitney; *P* <0.0001) (**K**). DNA is in blue in A, B, C, D, E, F, G and H. STAT-GFP brightness was increased in E’ and G’ to better visualise midguts. Scale bars in A, B, C, D, E, F, G and H are 50μm.

Together, our data indicate that ISCs sense midgut damage using Draper-Src-Shark signalling, which promotes ISC proliferation through STAT activation. Furthermore, Draper-Src-Shark signalling in ISCs, EBs, or in both progenitors cell non-autonomously activates STAT in the VM upon midgut damage.

### ISC Draper-Src-Shark signalling detects epithelial apoptosis after damage

Having shown that damage-induced Draper-Src-Shark signalling has an essential role in the regenerative response to midgut damage, we set out to identify the signal detected by Draper upon damage. Inhibition of apoptosis is known to partially block ISC proliferation that occurs after injury due to DNA damage ^51^. Thus, we tested whether inhibiting apoptosis in ECs blocks ISC proliferation after *P.e.* infection by expressing the apoptosis inhibitor p35 with an EC-specific driver, *mex-GAL4 tubGAL80*^ts^ (*mex*^ts^). We found that p35 expression in ECs indeed partially blocked ISC proliferation by 40.1% in *P.e.*-infected midguts compared to control infected midguts (Fig. 5A), suggesting that some of the infected ECs were undergoing apoptosis.

Draper recognises apoptotic cells by detecting phosphatidylserine on the outer leaf of the plasma membrane ^38, 43, 44^. Furthermore, Draper-Src-Shark signalling in glia can promote phagocytosis of axonal debris and apoptotic neuronal corpses after adult ORN injury ^40^. Thus, we tested whether progenitors with active Draper-Src-Shark signalling can engulf damaged or dying midgut epithelial cells. We marked ECs with nuclear ß-galactosidase (ß-gal) using *Myo1A-lacZ*. To test if apoptotic ECs were engulfed by midgut progenitors, we infected flies carrying *Myo1A-lacZ* with *P.e.* and determined if ß-gal from dying ECs could be detected within midgut progenitors. We found stable nuclear ß-gal expression in ECs after *P.e.* infection (Fig. 5D, E-E’; 5F, G-G”). However, we did not find nuclear ß-gal in Arm^high+^ Pros^-^ progenitors, indicating that *Myo1A* expression is not induced in progenitors after midgut damage (Fig. 5B, 5D, E-E’; 5F, G-G”). We then determined whether ß-gal from neighbouring ECs can be detected within Arm^high+^ Pros^-^ progenitors of control and *P.e.*-infected midguts. We found that 13.4% (mean) of progenitors within infected midguts were positive for ß-gal in their cytoplasm (Fig. 5C, 5F, G-G’’). This was similar to progenitors within uninfected control midguts (11.3%, mean) (Fig. 5C, 5D, E-E’’), indicating that dying ECs are not engulfed by progenitors. In support, we also did not observe GFP expressed in ECs using *mex*^ts^ within Arm^high+^ Pros^-^ progenitors, neither in control uninfected midguts (Supplementary Fig. 3D, E-E’’) nor in *P.e.*-infected midguts (Supplementary Fig. 3F, G-G’’). Together, these data indicate that progenitors use Draper-Src-Shark signalling to recognise apoptotic damaged cells but not engulf them.

In summary, we have here uncovered the mechanism by which *Drosophila* midgut progenitors recognise damage and translate this into increased STAT activity within progenitors and the visceral muscle to ensure regeneration (Figure 7). We propose that upon midgut damage, the immunoreceptor Draper on progenitors detects exposed phosphatidylserine on the plasma membrane of apoptotic enterocytes. This allows cortically-localised activated Src to phosphorylate the ITAMs on Draper. Subsequently, Shark binds to the phosphorylated ITAMs within Draper, allowing it to be further activated by Src through phosphorylation. Together, Draper-Src-Shark signalling in progenitors stimulates STAT activity within ISCs and EBs. Moreover, Draper-Src-Shark signalling in progenitors activates STAT in the visceral muscle, indicating that progenitors can shape the regenerative niche after tissue damage. Collectively, the increase of Stat activity within ISCs and the visceral muscle boosts ISC proliferation to promote regeneration.

**Figure 7.**
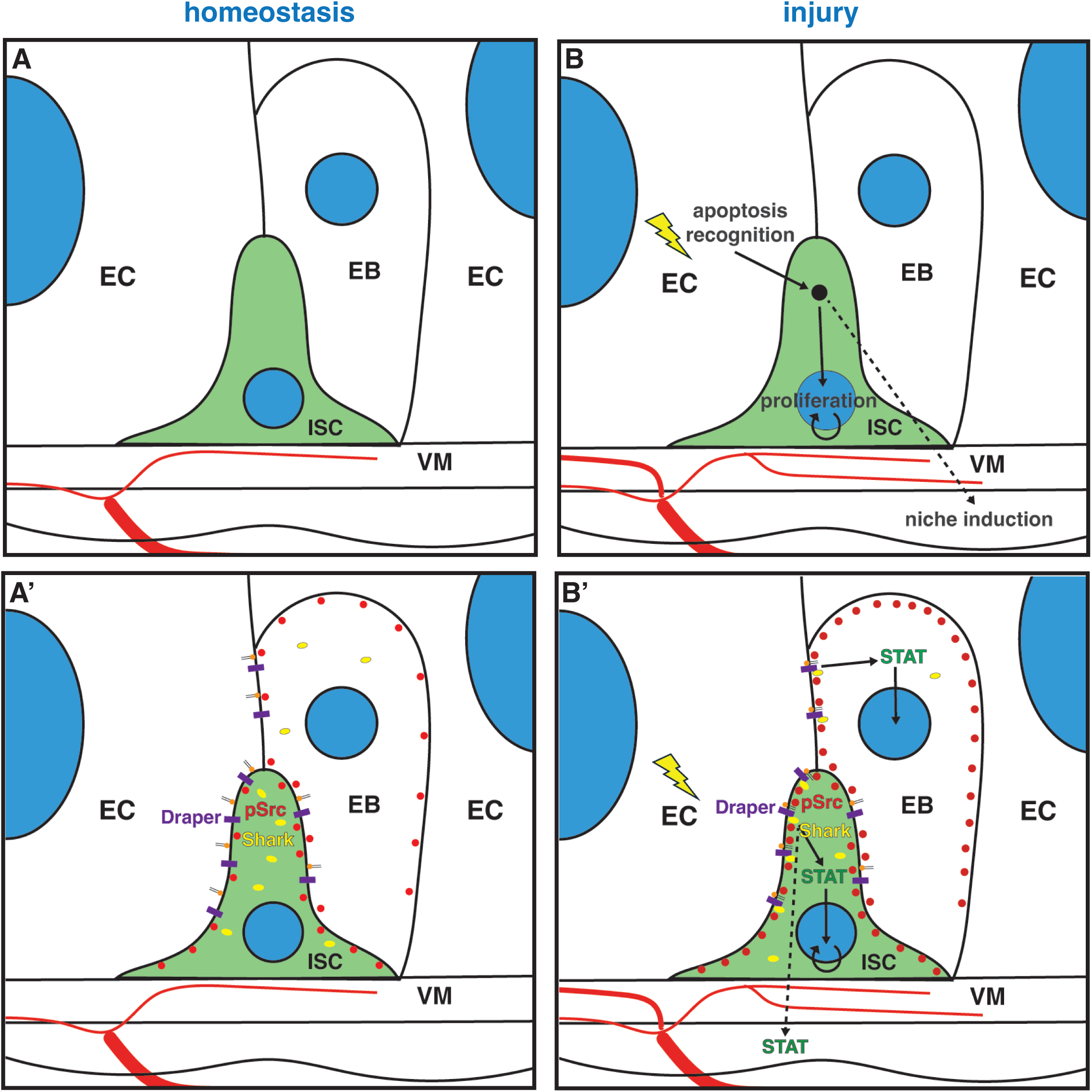
Damage recognition by Draper-Src-Shark-STAT signalling in intestinal stem cells promotes adult *Drosophila* midgut regeneration. Under homeostasis, a low level of phosphorylated Src is cortically localised within midgut progenitors (ISCs and EBs). The immunoreceptor Draper does not detect phosphatidylserine in undamaged cells and cannot associate with the Spleen tyrosine kinase homologue, Shark (**A, A’**). After injury, apoptosis is recognised by ISCs, which stimulates their proliferation and shapes the regenerative niche (**B**). Draper in progenitors detects exposed phosphotidylserine on dying enterocytes (ECs) after midgut damage, and a higher level of phosphorylated Src accumulates cortically in progenitors. Activated Src phosphorylates the immunoreceptor tyrosine-based activation motifs (ITAMs) on Draper. Shark binds to the phosphorylated ITAMs within Draper, allowing it to be further activated by Src through phosphorylation. Draper-Src-Shark signalling in progenitors activates STAT both in progenitors and within the visceral muscle (VM) (**B’**).

## Discussion

It is not well understood how Src signalling promotes ISC proliferation in both *Drosophila* and mammalian intestinal stem cells ^27^. We found that Src42A signalling is required in progenitors to respond to both pathogenic bacterial infection (*P.e.*) and detergent stress (SDS) (Fig. 1D, E). This suggests that Src42A in ISCs senses tissue damage rather than pathogenic bacteria themselves, possibly through the recognition of pathogen-associated molecular patterns (PAMPs). It has been previously shown that EBs may sense tissue damage after *P.e.* infection and stimulate ISC proliferation by producing Wingless (Wg) that activates Src in progenitors ^27^. Further, Ret overexpression in progenitors activates Src, which boosts their Wg levels and thus ISC proliferation ^34^. Ret is also required for ISC proliferation after damage ^34^, presumably by activating Src and bolstering Wg signalling. Whilst Wg and Ret signalling in progenitors after tissue damage promote ISC proliferation through Src activation, it was not clear whether Src activity was consequential in ISCs or EBs to induce ISC proliferation after tissue damage.

The progenitor population consists of three cell types, ISCs, EBs and EEps. We found that Src42A signalling is required substantially more in ISCs than in Su(H)^+^ EBs to promote ISC proliferation after midgut damage (Fig. 1F, G). As we used an EB-specific driver for EBs that have received a strong Notch signal (Su(H)^+^), it is possible that Src42A activity is still important in early EBs with low Notch signalling (Su(H)^-^) or in EEps for midgut regeneration. However, the odds that Src42A is vital in these cell types to regulate ISC proliferation after midgut damage is low as early EBs and EEps make up only about 5-10% of the *esg^+^* progenitor population. Whilst Wingless and Ret signalling have been shown to promote ISC proliferation via Src activation in progenitors, our work pinpoints the importance of Src42A activity in ISCs for their response to tissue damage.

Src42A has been shown to require EGFR/ERK and JAK-STAT signalling to promote ISC proliferation after midgut damage ^27, 33^. While STAT is crucial for Src42A to promote ISC proliferation, the Domeless receptor is not required ^27, 33^. This further suggests that cytokines (Upd1-3) are not required for Src42A to promote STAT activity and ISC proliferation. Thus, the mechanism by which Src42A regulates STAT activity was unknown.

Src family kinases are known to phosphorylate tyrosines within ITAM domains of immune receptors, thus allowing Syk kinases to bind to ITAMs through their SH2 domains ^36, 37^. Syk binding to ITAMs allows for its full activation by further phosphorylation by Src family kinases ^36, 37^. Indeed, we found activated Src42A to accumulate cortically in progenitors after *P.e.* infection (Fig. 1 A-C), likely allowing Src42A to phosphorylate the ITAM-containing immunoreceptor, Draper, and the Syk-homologue, Shark. In support, we found that Draper and Shark are both required in ISCs to stimulate their proliferation upon midgut damage (Fig. 3B and Fig. 4B). Draper is known to detect apoptotic cells through its binding to phosphatidylserine, which relocalises to the extracellular side of the plasma membrane during apoptosis ^38, 44^. Phosphatidylserine is normally found on the intracellular side of the plasma membrane within living cells. Draper signalling is utilised by both professional phagocytic hemocytes and non-professional phagocytes (glia, epithelia cells) to recognise apoptosis and clear apoptotic corpses through engulfment ^38–40^. We indeed found that EC apoptosis after *P.e.* infection is required for ISC proliferation (Fig. 5A). Additionally, we found that Draper is required in ISCs for their proliferation after *P.e.-*induced damage (Fig. 4B), suggesting that Draper on ISCs may recognise apoptotic midgut epithelial cells through phosphatidylserine recognition and engulf them. Interestingly, zebrafish epithelial stem cells and mouse hair follicle stem cells engulf apoptotic corpses ^52, 53^. Furthermore, the engulfment of dying epidermal cells by zebrafish epithelial stem cells promotes their proliferation to maintain tissue homeostasis ^53^. Nevertheless, we did not detect engulfment of apoptotic midgut ECs by progenitors during *P.e.* infection (Fig. 5C; D, E-E”; F, G-G’’; Supplementary Fig. 3D, E-E’’; F, G-G”). This could be due to their much larger size compared to ISCs or because apoptotic cells in the midgut extrude apically into the lumen ^54, 55^. Together, these data suggest that Draper on ISCs may sense apoptotic midgut epithelial cells but not engulf them. The Draper homologue in humans is also required in muscle satellite cells for skeletal muscle regeneration ^45^; however, how MEGF10 promotes regeneration is not understood. It would be intriguing to examine whether MEGF10 acts as a sensor for apoptotic cells on various stem cells after tissue injury, suggesting a conserved mechanism.

When Draper binds to phosphatidylserine on dying cells, it is phosphorylated by Src42A, which enables Shark binding and activation ^40, 43, 44^. We found that Shark is required for constitutive active Src42A to promote ISC proliferation (Fig. 3C). However, it was still unclear how Draper-Src-Shark signalling in ISCs activates ISC proliferation. Interestingly, STAT is activated by Draper-Src42A-Shark signalling in glial cells after injury to adult *Drosophila* ORN axons ^42^. Furthermore, STAT is required for ISC proliferation after pathogenic bacterial infection ^18, 26^. Moreover, STAT is thought to be exclusively activated by niche-derived cytokine Upd1-3 signalling ^20, 35, 56^. Upon pathogenic bacterial infection, Upd1 is produced by progenitors and early ECs; Upd2, by EBs and ECs and Upd3, by ECs and Malpighian tubules ^18–21, 57^. The importance of niche cytokines for ISC proliferation after midgut damage is demonstrated by the near-complete abolition of ISC proliferation in *upd2* and *upd3* double mutants after *P.e*. infection ^58^. While these data suggest a strong activation of STAT by niche-produced cytokines, we found that Draper-Src42A-Shark signalling in ISCs is required for full STAT activation in progenitors and the visceral muscle after *P.e.* infection (Figs. 2-4, 6). Indeed, we found a 41.4%, 62.6% and 58.7% decrease in STAT activation in *Src42A-*, *shark*- and *draper*-depleted progenitors, respectively, in *P.e.-*infected midguts (Supplementary Fig. 1C, 2C, 3C). These data suggest a role for Draper-Src-Shark signalling to regulate STAT activity in EBs or in ISCs to cell non-autonomously regulate STAT activity in EBs (Fig. 7). Furthermore, when *draper*, *shark* and *Src42A* were depleted from progenitors, we found that STAT activity diminished by 27.0%, 57.9% and 37.1%, respectively, in the VM of infected midguts (Fig. 6I-K). This suggests that ISCs, EBs or both ISCs and EBs can non-autonomously regulate the regenerative niche after midgut damage, possibly as a fail-safe mechanism.

How Stat3 is activated in the mouse intestinal epithelium after damage is not fully understood. Similar to *Drosophila*, Stat3 is thought to be largely activated by cytokines—IL-6 and IL-10 are produced by macrophages and IL-22 is produced by type-3 innate lymphoid (ILC3) cells ^5, 6^. Nevertheless, Stat3 may also be partly regulated by ISC intrinsic factors that sense intestinal damage. This is supported by the fact that Src-STAT3 signalling in the crypt containing ISCs and transit amplifying progenitors is critical for mouse intestinal epithelial regeneration after DNA damage by γ-irradiation ^27^. Furthermore, Src is required for Stat3 activation after γ-irradiation, indicating that epithelial Src activity is crucial for Stat3 activation after injury ^27^. Nevertheless, it is not clear how Src regulates Stat3 after injury. Whether Src in intestinal progenitors promotes damage sensing and activates Stat3 through MEGF10 or Syk as in our model remains to be determined.

Src is also activated in non-invasive human adenomas ^27^. Thus, it would be interesting to examine whether stem-like tumour cells utilise Src signalling to detect apoptotic cells within tumours or normal neighbouring tissue to fuel their growth. Alternatively, stem-like tumour cells could utilise Src signalling to promote tumour regrowth after chemotherapy or radiation treatment. In summary, our work increases our understanding of how stem cells recognise damage, couple it to their proliferation and influence their surrounding niche to support tissue regeneration. We hope that this knowledge will aid in the development of treatments for epithelial regeneration after injury, pathogenic infection, damage caused by inflammatory disease and cancer therapies.

## Materials and Methods

### Fly stocks

The following fly stocks were used: *w^1118^*, *w^1118^*; *10X-STAT-DGFP* ^49^, w; *esgGAL4; tubGAL80*^ts^ *UAS-GFP* (*esg*^ts^*GFP*), *w; esgGAL4 UAS-mRFP; tubGAL80*^ts^ (*esg*^ts^*mRFP*), *esgGAL4 UAS-2X EYFP; Su(H)Gbe-GAL80, tub-GAL80*^ts^ (*esg*^ts^ *Su(H)GAL80*), *w; Su(H)Gbe-GAL4; tub-GAL80*^ts^ *UAS-GFP* (*SuH*^ts^); *mex-GAL4; tubGAl80*^ts^ (*mex*^ts^), *UAS-p35* and *Myo1A-lacZ* ^18^; *UAS-Src42A^RNAi^* (2)(26019GD), *UAS-shark^RNAi^* (2) (105706KK)^59^, *UAS-draper^RNAi^* (1)(4833GD) and *draper^RNAi^* (2)(27086GD) from the Vienna *Drosophila* Stock Center (VDRC) and *UAS-Src42A^CA^*, *UAS-Src42A^RNAi^* (1)(HMC04138)^60^, *UAS-shark^RNAi^* (1)(HMC04146)^61^, *yv*; *UAS-GFP*, *yv*; *UAS-mCherry^RNAi^* from the Bloomington *Drosophila* Stock Center (BDSC).

### *Drosophila* genetics

Flies raised at 18°C were shifted to 29°C to induce UAS transgene expression. Experiments were performed using 5-10 day old, adult, female *Drosophila melanogaster*. Typically, between 10-20 flies were used per experiment. Flies were selected by their genotype and then randomly chosen to be used in an experiment.

### Stress experiments

*Pseudomonas entomophila* (*P.e.*) infection: flies were treated for 18 hours with food laced with 500μl of 5% sucrose or 20x *P.e.* resuspended in 5% sucrose from an 18-hour culture grown at 30°C, 130 RPM. Detergent exposure: 500μl of 0.2% sodium dodecyl sulfate (SDS) or H20 was mixed into the food.

### Histology

Following fixation in 8% formaldehyde for 30-60 minutes, midguts were washed in PBS, 0.1% Triton X-100, blocked in PBS, 0.1% Triton X-100, 1% bovine serum albumin (BSA), 2% normal goat serum (NGS), and stained in blocking solution with rabbit polyclonal anti-phospho Ser 10 histone 3 (Merck, 06-570; 1:1000), mouse monoclonal anti-phospho Ser 10 histone 3 (3H10) (Sigma Aldrich, 05-806; 1:1000), rabbit polyclonal anti-phosphorylated human Src Y419 (Invitrogen, 44-660G; 1:100), mouse anti-Armadillo (DHSB, N2 7A1; 1:50), mouse anti-Prospero (DHSB, MR1A; 1:50), chicken anti-ß-galactosidase (Abcam, ab9361; 1:1000), chicken polyclonal anti-GFP (Life Technologies, A10262; 1:1000) or rabbit polyclonal anti-RFP (Thermo Fisher Scientific, R10367; 1:1000). Midguts were then washed in PBS 0.1% Triton X-100, stained in PBS, 0.3% Triton X-100, 0.1% BSA with Alexa Fluor-conjugated secondary antibodies (Life Technologies, 1:1000) and Hoechst 33258 (Life Technologies) and mounted in Vectashield (Vector Laboratories). For anti-phospho-Src, midguts were fixed in 8% formaldehyde containing PhosSTOP.

### Microscopy

Samples were analysed using a Leica DM16000 and Leica SP5 microscope. Images were processed with ImageJ (NIH) and Adobe Photoshop. Confocal images are presented as maximum intensity projections of images. A z-stack for each experiment was acquired with the same laser intensity and gain.

The number of mitoses per midgut was determined by counting the number of phospho-Ser 10 histone 3-positive cells from whole female midguts. The mean number of mitoses per midgut and 95% confidence interval are presented for each genotype and treatment.

### Image quantitation

*Phospho-Src in progenitors:* average intensity projections of 12 slices from 0.25μm step z-stacks of the basal layer containing progenitors were generated for the *esg*>GFP^+^ and Armadillo channels and merged using ImageJ. Similarly, average intensity projections of 12 slices were generated for the pSrc channel using ImageJ, and rolling ball (10 pixel radius) background subtraction was applied prior to quantitation. *esg*>GFP^+^ cells for each condition were randomly selected and the inner side of the plasma membrane was traced (line thickness of 3) with the ImageJ freehand selection tool using the Armadillo signal as a guide. These selections were restored onto pSrc projections, and the raw integrated density was measured along the line. The raw integrated density was then multiplied by 12 and divided by the line length to obtain the normalised fluorescence intensity of pSrc at the membrane per cell.

*Stat-GFP in progenitors:* sum projections of 10 slices from 0.5 μm z-stacks of the basal layer containing progenitors (high Arm^+^) were generated for the Arm and 10X STAT-DGFP channels using ImageJ. The Arm signal within high Arm^+^ cells was traced using the freehand selection tool, and the selection was restored onto 10X Stat-DGFP projections to measure the raw integrated density within the cell. The raw integrated density for each progenitor was then divided by the area of each progenitor to determine the normalised fluorescence intensity of 10X STAT-DGFP per cell.

*Stat-GFP in visceral muscle:* sum projections were generated from 15 slices of 0.13μm z-stacks for the DNA (Hoechst) and 10X STAT-DGFP channels using ImageJ. The visceral muscle nuclei were traced using the freehand selection tool. These selections were then restored onto the 10X STAT-DGFP channel to measure the raw integrated density for 10X STAT-DGFP within nuclei. The raw integrated density for each nucleus was then divided by the area of each nucleus to determine the normalised fluorescence intensity of 10X STAT-DGFP per nucleus.

*Nuclear and cytoplasmic* β*-gal in progenitors:* maximal intensity projections of 30 slices of 0.25μm z-stacks were generated for the Arm and β-gal channels in ImageJ and merged in Adobe Photoshop. All Arm^high+^ Pros^-^ cells were then manually assessed for the expression of nuclear β-gal or for increased amounts of cytoplasmic β-gal signal that could indicate enterocyte engulfment. Approximately 100-130 cells per midgut were examined.

*esg^+^ progenitor cell /total cell ratio:* maximal intensity projections of 10 slices of 1 μm z-stacks were generated for the esg>GFP and DNA (Hoechst) channels in ImageJ. The total number of *esg^+^* cells with small nuclei and the total number of nuclei were then counted. The number of esg^+^ cells was then divided by the total number of nuclei to obtain the ratio *esg^+^* cells/ total cells.

### Statistical analysis

Statistical analyses were performed using Graphpad Prism 10. A two-sided unpaired t-test or Mann-Whitney test (for non-Gaussian data) were applied to determine statistical significance. The significance level is indicated by an * for p<0.05, ** for p<0.01, *** p<0.001, and **** for p<0.0001, and ns, for not significant, p>0.05.

## Acknowledgements

We thank the Bloomington *Drosophila* Stock Center and the Vienna *Drosophila* RNAi Center for fly stocks and the Developmental Studies Hybridoma Bank (DHSB) for antibodies.

This research was funded in whole by the Wellcome Trust and Royal Society (Grant Number 220198/Z/20/Z) to P.H.P.. For the purpose of open access, the author has applied a CC BY public copyright licence to any Author Accepted Manuscript version arising from this submission.

## Author contributions

P.H.P. conceived, developed and supervised the project. P.H.P. designed the experiments, and P.H.P., M.L., R.W. and M.S. performed and analysed the experiments. M.L. contributed to Fig. 1C-E, 1G, Fig. 3-5, Fig. 6A-J, Supplementary Figs. 1A, 2A, 2C, 3A, 3C-G”; R.W. contributed Fig. 1A-C, 1F, Supplementary Figs. 1B, D-G’, 2B and 3B; M.S. contributed Fig. 2, Fig. 6K, Supplementary Figs. 1C, 1H-K’. M.L., M.S. and P.H.P. prepared the figures and figure legends. P.H.P. wrote and edited the manuscript.

